# Olfactory inputs to appetite neurons in the hypothalamus

**DOI:** 10.1101/2023.02.28.530282

**Authors:** Donghui Kuang, Naresh K. Hanchate, Chia-Ying Lee, Ashley Heck, Xiaolan Ye, Michidsaran Erdenebileg, Linda B. Buck

## Abstract

The sense of smell has potent effects on appetite, but the underlying neural pathways remain undefined. Here we investigated how olfactory signals reach two subsets of appetite-linked neurons in the hypothalamic arcuate nucleus: AgRP (agouti-related peptide) neurons, which stimulate appetite, and POMC (pro-opiomelanocortin) neurons, which suppress it. Using polysynaptic viral tracing, we show that AgRP and POMC neurons receive indirect input from partially overlapping but distinct areas of the olfactory cortex, indicating that they process different sets of olfactory information. We also identify different complements of neurons directly upstream of AgRP and POMC neurons that can relay olfactory cortical signals to the appetite neurons. Single cell transcriptomics shows heterogeneous expression of neuromodulator receptors among AgRP neurons, suggesting variations in the signals they receive. Integrated viral tracing and RNA localization further reveals selected brain areas where upstream neurons express cognate receptor ligands. Together, these findings outline multiple pathways by which distinct olfactory and modulatory signals are differentially routed to neurons that promote versus inhibit appetite.

## Introduction

The regulation of appetite and food intake is essential for survival and is conserved across the animal kingdom. In mice and other mammals, the arcuate nucleus of the hypothalamus (ARC) contains two subsets of neurons with opposite roles in appetite regulation: AgRP neurons, which promote appetite, and POMC neurons, which inhibit it. AgRP neurons express agouti-related peptide, while POMC neurons express pro-opiomelanocortin (POMC)^1,2^.

AgRP neurons are activated when animals are calorically deficient^3–5^. Optogenetic or chemogenetic activation of these neurons stimulates voracious eating and other hunger-related behaviors^6–8^. Activated AgRP neurons can also induce lipogenesis, fat mass accumulation, and altered substrate utilization^9^. In contrast, POMC neurons are stimulated when energy is available^5^, and optogenetic activation of POMC neurons inhibits feeding even in fasted animals^10^.

The smell and sight of food dramatically alters AgRP and POMC neuron activity^11–13^. Within seconds before food is consumed, the activity of AgRP neurons is inhibited while POMC neurons are stimulated. These effects appear to arise from learned associations between sensory food cues and the caloric or nutritional value of the food^11,14^.

How is the smell of food conveyed to AgRP and POMC neurons to elicit these effects on appetite? In the olfactory system, odor signals travel from the nasal olfactory epithelium through the olfactory bulb to the olfactory cortex, which transmits information to multiple other brain areas^15,16^. The olfactory cortex comprises a number of anatomically distinct areas whose respective functions are poorly understood^17–19^. Whether one or more of these areas is involved in appetite is unknown.

Previous studies used monosynaptic rabies virus to map the locations of brain neurons presynaptic to AgRP or POMC neurons^20,21^. Those studies showed neurons presynaptic to AgRP or POMC neurons in multiple brain areas, but not the olfactory cortex. Those results suggested that odor signals might be conveyed from the olfactory cortex to AgRP and POMC neurons via indirect routes that cannot be detected using a monosynaptic rabies virus.

To investigate this idea, we infected AgRP or POMC neurons with PRVB177, a conditional Pseudorabies virus that travels retrogradely across multiple synapses after infecting neurons expressing Cre recombinase^22^. These experiments revealed infected neurons likely to be two synapses upstream of AgRP and POMC neurons in the olfactory cortex (OC), confirming that both receive odor signals indirectly from the OC. However, infected neurons were seen in some OC areas but not others, suggesting that only some OC areas provide signals to the appetite neurons. In addition, OC areas upstream of AgRP and POMC neurons overlapped only partially, further suggesting that AgRP and POMC neurons receive different sets of olfactory information.

Neurons directly upstream of AgRP or POMC neurons were seen in multiple brain areas. These areas provide potential routes by which information could be relayed from the OC to the appetite neurons to affect their activity and thus modulate appetite.

Neurons upstream of AgRP and POMC neurons were also present in two amygdala areas that receive signals from the vomeronasal organ, an accessory olfactory structure that detects social cues^23^. This finding raises the possibility that social cues could influence appetite.

To investigate signaling mechanisms that could relay olfactory information to AgRP neurons, we employed single cell RNA sequencing to identify AgRP neuron receptors for neuromodulators. We then used viral tracing combined with RNA in situ hybridization to locate upstream neurons expressing cognate neuromodulators. These experiments uncovered numerous neuromodulator receptors differentially expressed by AgRP neurons. They also revealed that cognate ligands of those receptors can be selectively expressed by upstream neurons in specific brain areas.

Together, these studies uncover areas of olfactory cortex that convey odor signals to AgRP and POMC neurons and non-olfactory brain areas with the potential to relay those signals from the olfactory cortex to AgRP and POMC neurons. These studies also unravel signaling mechanisms that can modulate AgRP neurons. They show that single AgRP neurons differentially express diverse receptors for neuromodulators and that ligands of those receptors are selectively expressed in neurons upstream of AgRP neurons in specific brain areas. These findings provide a foundation for molecular-genetic studies to further explore the transmission of olfactory information to AgRP and POMC appetite neurons and signaling circuits and mechanisms underlying the modulation of appetite.

## Results

### Appetite neurons receive indirect input from the olfactory cortex

To investigate whether AgRP and POMC neurons receive input from higher olfactory areas, we utilized PRVB177, a Cre-dependent Pseudorabies virus (Fig. 1). PRVB177 has an irreversible Cre recombinase-dependent hemagglutinin tagged thymidine kinase (HA-TK), which allows it to propagate and travel retrogradely across multiple synapses^22^.

**Fig. 1.**
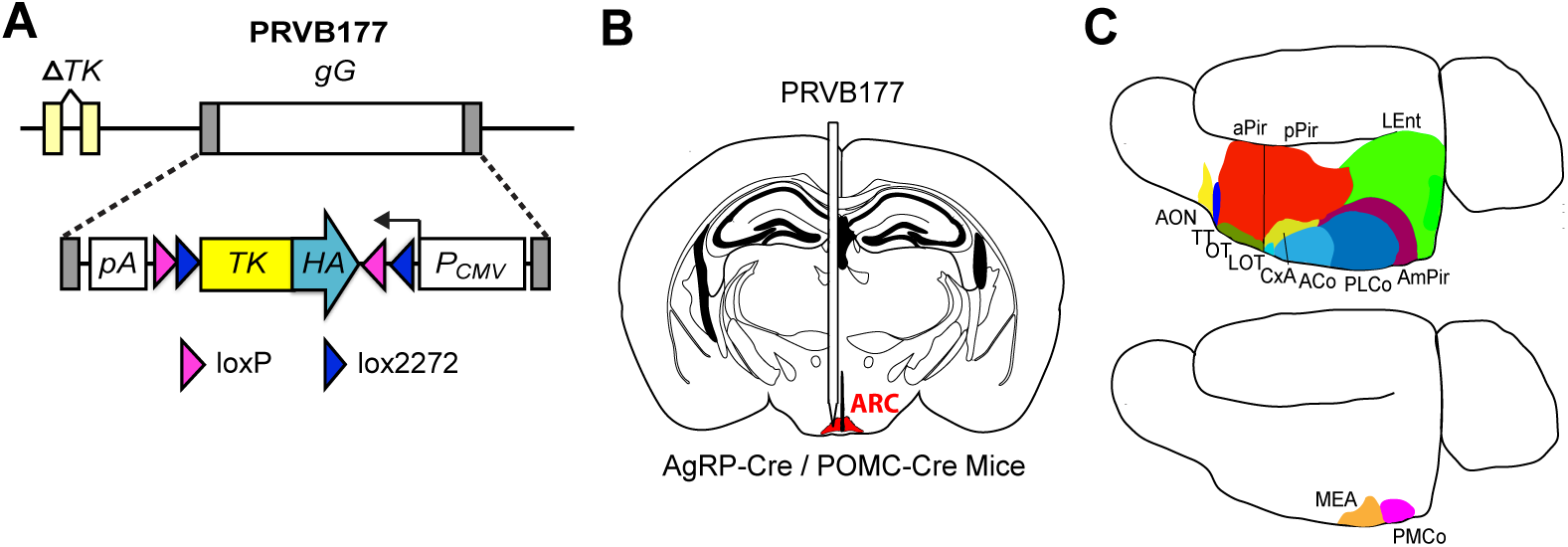
Strategy to assess input to ARC AgRP and POMC neurons from higher olfactory areas. **(A)** Cre recombinase-driven irreversible expression of TK-HA from PRVB177 allows viral replication and synaptic spread. *gG*, *gG* locus; *PCMV*, cytomegalovirus promoter; *pA*, polyadenylation signal. **(B)** Injection of PRVB177 into the ARC (indicated in red) of AgRP-Cre or POMC-Cre mice. **(C)** Immunostaining for PRV (HA) in brain sections to identify infected neurons within olfactory cortex (above) or vomeronasal amygdala (below).

PRVB177 was injected into the arcuate nucleus (ARC) of mice expressing Cre in AgRP or POMC neurons (AgRP-Cre or POMC-Cre mice) (Fig. 1). To examine the locations of upstream PRV-infected neurons, brain sections were then immunostained for HA. This method was previously used in CRH-Cre mice to examine neurons upstream of hypothalamic neurons expressing CRH (corticotropin releasing hormone)^22^.

PRV-infected (HA+) neurons were first detected outside the ARC on day 3 post-injection (d3pi), suggesting that these neurons are directly upstream of AgRP or POMC neurons. This is consistent with studies using CRH-Cre mice, which showed that virus-infected neurons seen on d3pi are one synapse upstream of (presynaptic to) infected Cre-expressing neurons whereas those first seen on d4pi are likely two synapses upstream^22^.

The olfactory cortex receives odor signals generated in the nose via a relay in the olfactory bulb. The OC comprises multiple anatomically distinct areas whose respective functions are poorly understood^17–19^ (Fig. 1). On d3pi, PRV+ neurons were observed in POMC-Cre mice in one OC area, the lateral olfactory tract (LOT). None were detected in the OC in AgRP-Cre mice (Fig. 2).

**Fig. 2.**
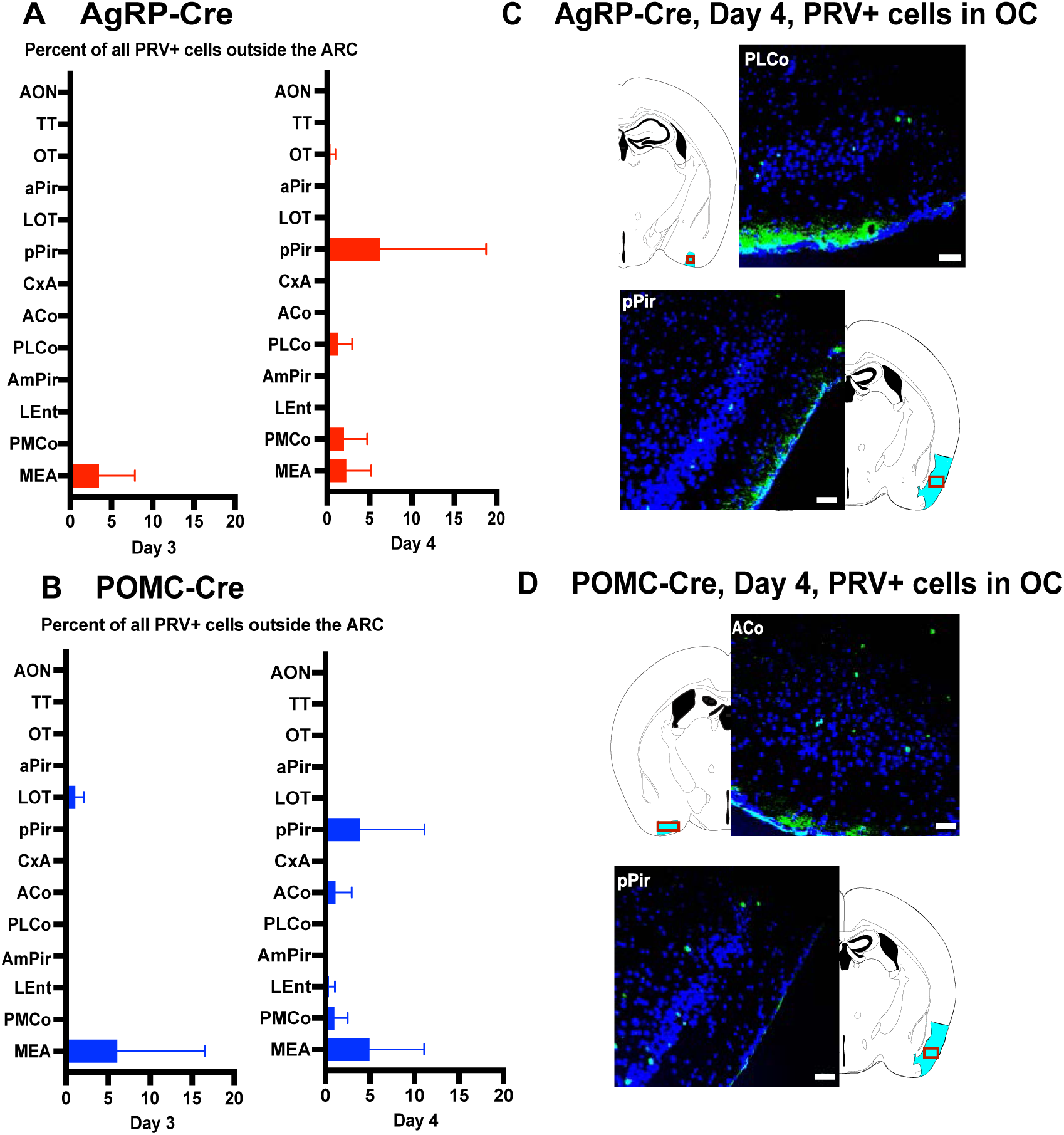
AgRP and POMC neurons receive input from higher olfactory areas. Mean percentages of PRV+ neurons outside ARC in areas of olfactory cortex or vomeronasal amygdala on day 3 or 4 after infection of AgRP (A) or POMC (B) neurons with PRVB177. Day 3: AgRP-Cre (*n* = 16); POMC-Cre (*n* = 4). Day 4: AgRP-Cre (*n* = 6); POMC-*Cre* (*n* = 4). Error bars indicate SEM. See Methods for full names of abbreviated brain areas. Photographs and diagrams of olfactory cortex sections with neurons immunostained for PRV on day 4 post-injection of AgRP-Cre (C) or POMC-Cre (D) mice. PRV+(HA+), green; DAPI counterstain, blue. Scale bars, 100 μm. Corresponding areas on diagrams are labeled with red rectangles in OC areas indicated in cyan.

In contrast, on d4pi, PRV+ neurons were seen in the OC of both AgRP-Cre and POMC-Cre mice (Fig. 2 and 3, Fig. S1). These neurons are likely to be two synapses upstream of the appetite neurons and thus provide indirect input to them. However, some OC areas contained PRV+ neurons whereas others did not. Of eleven different OC areas, five contained PRV+ neurons upstream of AgRP or POMC neurons, suggesting that only some OC areas provide signals to the appetite neurons.

**Fig. 3.**
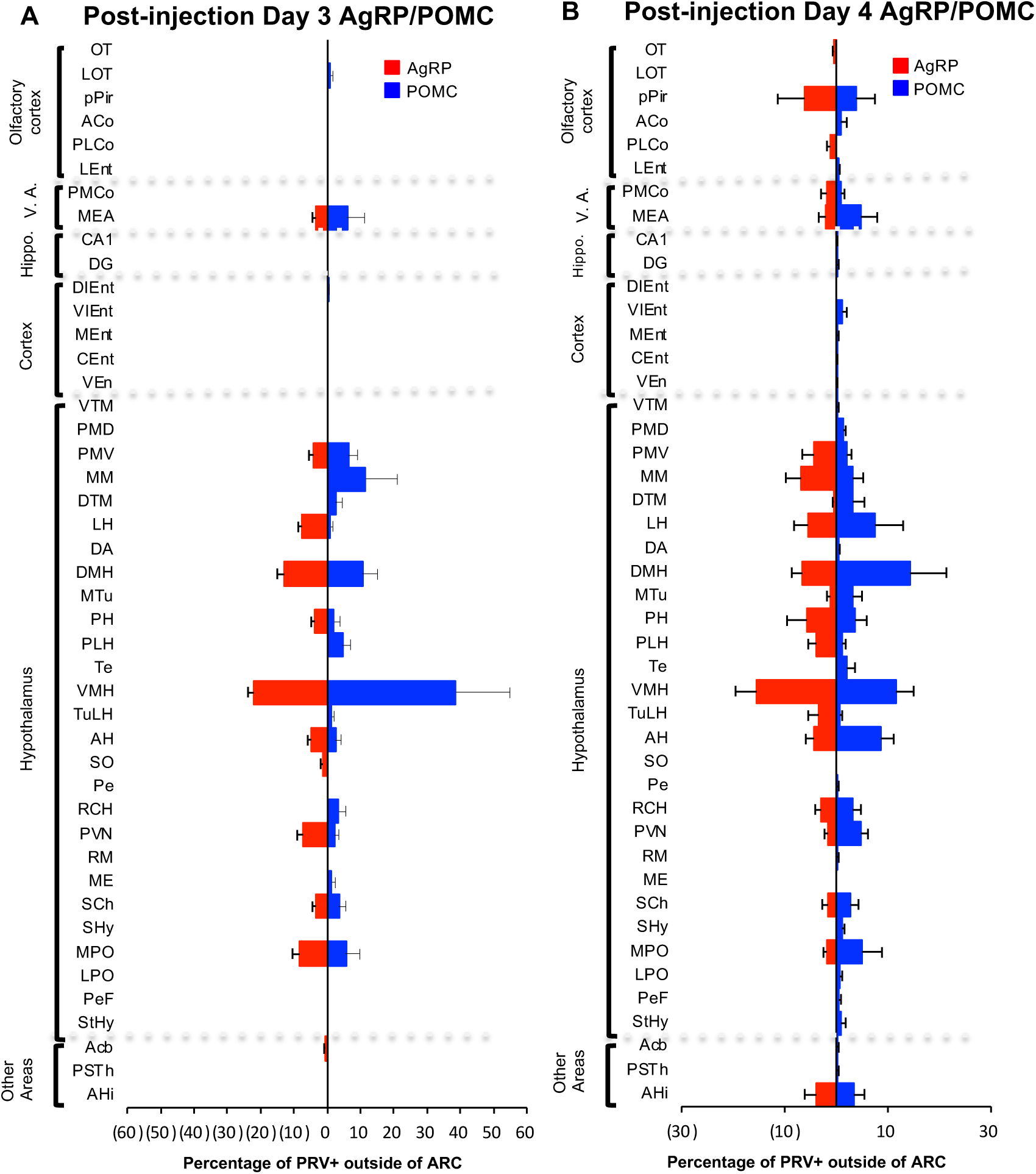
Brain areas with PRV+ neurons after infection of AgRP or POMC neurons. Bar graphs show the percentage of non-ARC PRV+ neurons in individual brain areas on days three (A) and four (B) post-infection of ARC AgRP (red) or POMC (blue) neurons. Sample sizes for days 3 and 4 post-injection as in Fig. 2. Error bars indicate SEM. V.A., vomeronasal amygdala; Hippo., hippocampus. See Methods for full names of abbreviated brain areas.

In addition, the OC areas with PRV+ neurons differed in AgRP-Cre versus POMC-Cre mice. In AgRP-Cre mice, PRV-infected neurons were identified in three OC areas: the olfactory tubercle (OT), posterior lateral cortical amygdala (PLCo), and posterior piriform cortex (pPir). The percentages of all PRV+ cells outside the ARC in these areas were 0.4 ± 0.3%, 1.2 ± 0.7%, and 6.1 ± 5.2%, respectively. In POMC-Cre mice, PRV+ neurons were seen in a different set of three OC areas: the lateral entorhinal cortex (LEnt), anterior cortical nucleus (ACo), and pPir. The percentages of all PRV+ cells external to the ARC in these three areas were 0.4 ± 0.3%, 1.0 ± 1.0%, and 3.8 ± 3.7%, respectively.

These results indicate that both AgRP and POMC appetite neurons receive information indirectly from the olfactory cortex. They suggest that one major OC area, the pPir contains neurons two synapses upstream of both AgRP and POMC neurons. They further suggest that there are also neurons two synapses upstream of AgRP neurons in two additional OC areas, the OT and the PLCo and neurons two synapses upstream of POMC neurons in two other OC areas, the ACo and the LEnt. Thus, while one OC area, pPir, provides indirect input to both subsets of appetite neurons, there are four additional OC areas that provide indirect input to only one or the other subset: the OT and PLCo only to AgRP neurons and the ACo and LEnt only to POMC neurons. Thus, AgRP and POMC neurons receive signals from partially overlapping areas of the OC and could thereby receive different sets of olfactory information.

Mice have an accessory olfactory structure in the nasal septum called the vomeronasal organ (VNO), which detects pheromones and other social cues^15,16,24^. PRV+ neurons were observed upstream of both AgRP and POMC neurons in the “vomeronasal amygdala” (VA), which receives signals derived from the VNO (Fig. 1, 2, and 3, Fig. S1). One VA area, the medial amygdala (MEA), contained PRV+ neurons upstream of both AgRP and POMC neurons on d3pi, suggesting that it provides direct input to the appetite neurons. The other, the posteromedial cortical amygdala (PMCo), showed PRV+ neurons upstream of both types of appetite neurons on d4pi, suggesting indirect inputs to the appetite neurons. These results suggest a means by which social cues detected in the VNO might influence appetite.

Together, these results indicate that the OC provides indirect input to both AgRP and POMC neurons. However, only some areas of the OC do this. Moreover, AgRP and POMC neurons receive input from partially overlapping but distinct OC areas, suggesting that they receive different complements of olfactory information.

### Appetite neurons receive direct input from multiple brain areas

The presence of OC neurons indirectly upstream of AgRP and POMC neurons implied that OC signals are relayed to the appetite neurons by their presynaptic partners in one or more other brain areas. These experiments identified PRV+ neurons directly upstream of AgRP or POMC neurons in multiple areas that could play this role. These PRV+ neurons were identified on d3pi, when the virus has crossed one synapse^22^ and are thus likely presynaptic to the appetite neurons.

On d3pi of AgRP neurons, PRV+ cells were identified in 11 non-olfactory (non-OC, non-VA) brain areas outside the ARC (Fig. 3, Fig. S1, S2, and S4). These areas were predominantly in the hypothalamus, a region that regulates endocrine functions and other basic functions, such as appetite, thirst, and instinctive behaviors^1,25–31^. The highest concentrations of PRV+ cells were observed in the ventromedial hypothalamus (VMH) and dorsal medial hypothalamus (DMH). PRV+ cells were also seen in the nucleus accumbens (Acb), which is linked to reward^32^.

On d3pi of POMC neurons, PRV+ neurons were detected in 16 non-olfactory brain areas (Fig. 3, Fig. S1, S3, and S4). As with AgRP neurons, these areas were primarily located within the hypothalamus, with high concentrations of PRV+ cells in both the VMH and DMH.

Nine non-olfactory brain areas exhibited PRV+ neurons 3 days post-infection of both AgRP and POMC neurons, suggesting that these regions could modulate both neuron types (Fig. 3, Fig. S1 and S4). It remains to be determined whether the neurons in these areas that project to AgRP and POMC neurons are identical or distinct. These findings are similar to those from studies employing a monosynaptic rabies virus to trace neurons presynaptic to AgRP and POMC neurons^20,21^.

By d4pi, additional non-olfactory brain areas showed PRV+ neurons upstream of AgRP or POMC neurons (Fig. 3, Fig S1). As in the OC, the new PRV-infected neurons identified on day 4 are likely to be two synapses upstream of AgRP or POMC neurons and transmit information to the appetite neurons indirectly via relays directly upstream of the appetite neurons.

These experiments showed eleven brain areas with neurons directly upstream of AgRP neurons and sixteen with neurons directly upstream of POMC neurons. The upstream neurons in one or more of those areas could relay signals from the OC to the appetite neurons. Most or all projection neurons in the OC are excitatory glutamatergic neurons. However, they could activate either excitatory or inhibitory neurons presynaptic to AgRP or POMC neurons and thereby either stimulate or suppress the appetite neurons.

### AgRP neurons express diverse receptors for neuromodulators

The above experiments traced neurons directly upstream of AgRP neurons to 11 non-olfactory brain areas, one or more of which could relay information from the OC to AgRP neurons. To explore signaling mechanisms that could relay olfactory information to AgRP neurons, our strategy was to catalog AgRP neuron receptors for neuromodulators and then pinpoint the sources of receptor ligands in upstream neurons (Fig. 4).

**Fig. 4.**
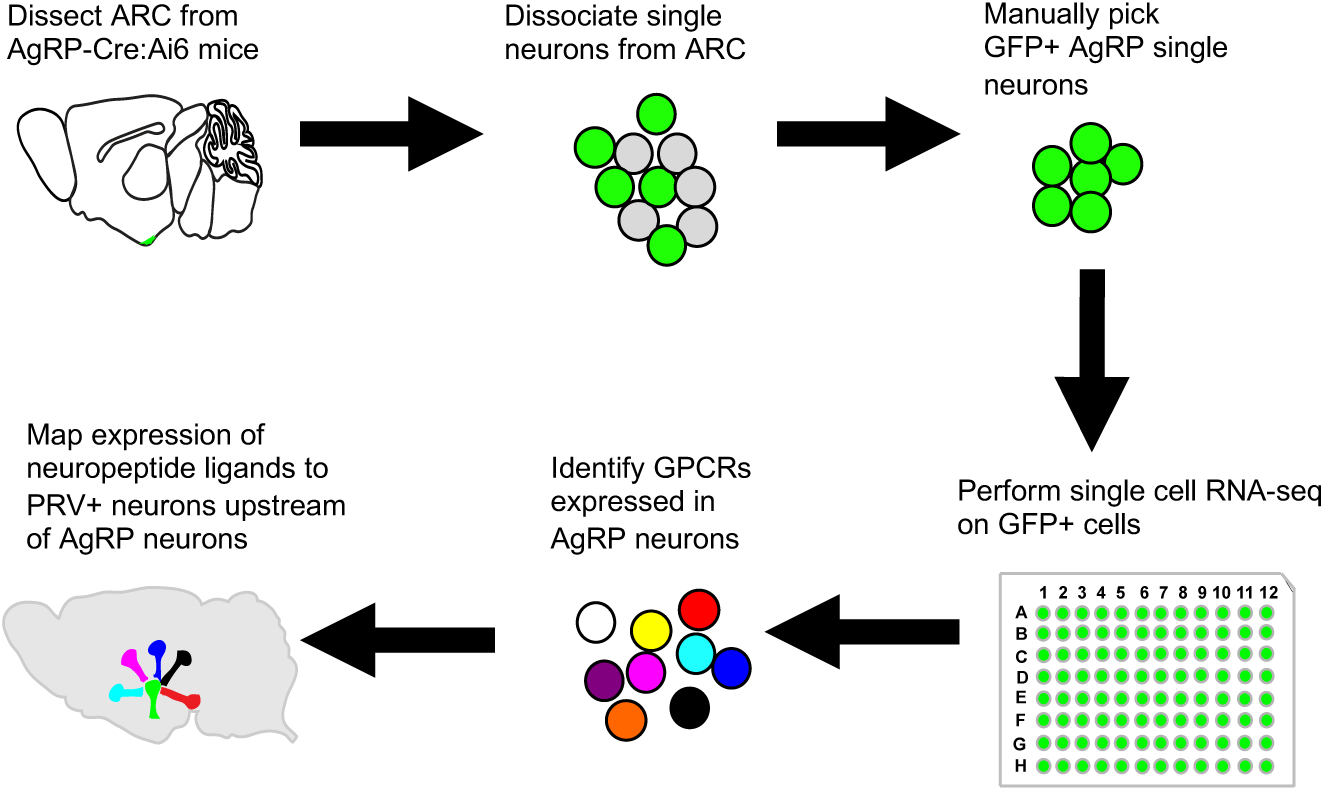
Identification of GPCRs expressed in AgRP neurons and upstream neurons expressing their ligands. The ARC was dissected from AgRP-Cre:Ai6 mice and dissociated into single cells. Single GFP+ AgRP neurons were manually isolated and subjected to single cell RNA-seq. Transcriptome data was used to identify GPCRs expressed in single AgRP neurons and those with known ligands determined. The expression of individual neuropeptide ligands in neurons directly upstream of AgRP neurons was determined by infecting AgRP neurons with PRVB177 and costaining brain sections for PRV and different neuropeptides.

To identify AgRP neuron receptors for neuromodulators, single cell transcriptome analysis was performed on AgRP neurons isolated from the arcuate nucleus. AgRP-IRES-Cre mice^33^ were crossed with Ai6 mice^34^, which have Cre-dependent expression of eGFP, to generate AgRP-Cre:Ai6 mice. Individual GFP+ cells were isolated from the ARC and subjected to single-cell RNA sequencing (scRNA-seq), employing established protocols to generate and analyze single-cell cDNA libraries^28,35,36^. On average, cells were sequenced at ∼6.5 million reads, with the sequenced reads mapped to the mouse genome (UCSC mm10, GENCODE M15). Gene expression was quantified, setting a threshold of 1 FPKM for expression in individual cells.

This analysis revealed a diverse array of G protein-coupled receptors (GPCRs) expressed by AgRP neurons (Fig. 5, Fig. S5)^37^. These included eight metabotropic receptors for the neurotransmitters glutamate, GABA, or ADP/ATP, six receptors for the biogenic amines epinephrine/norepinephrine, histamine, or serotonin, 23 receptors for neuropeptides, and four for other signaling molecules.

**Fig. 5.**
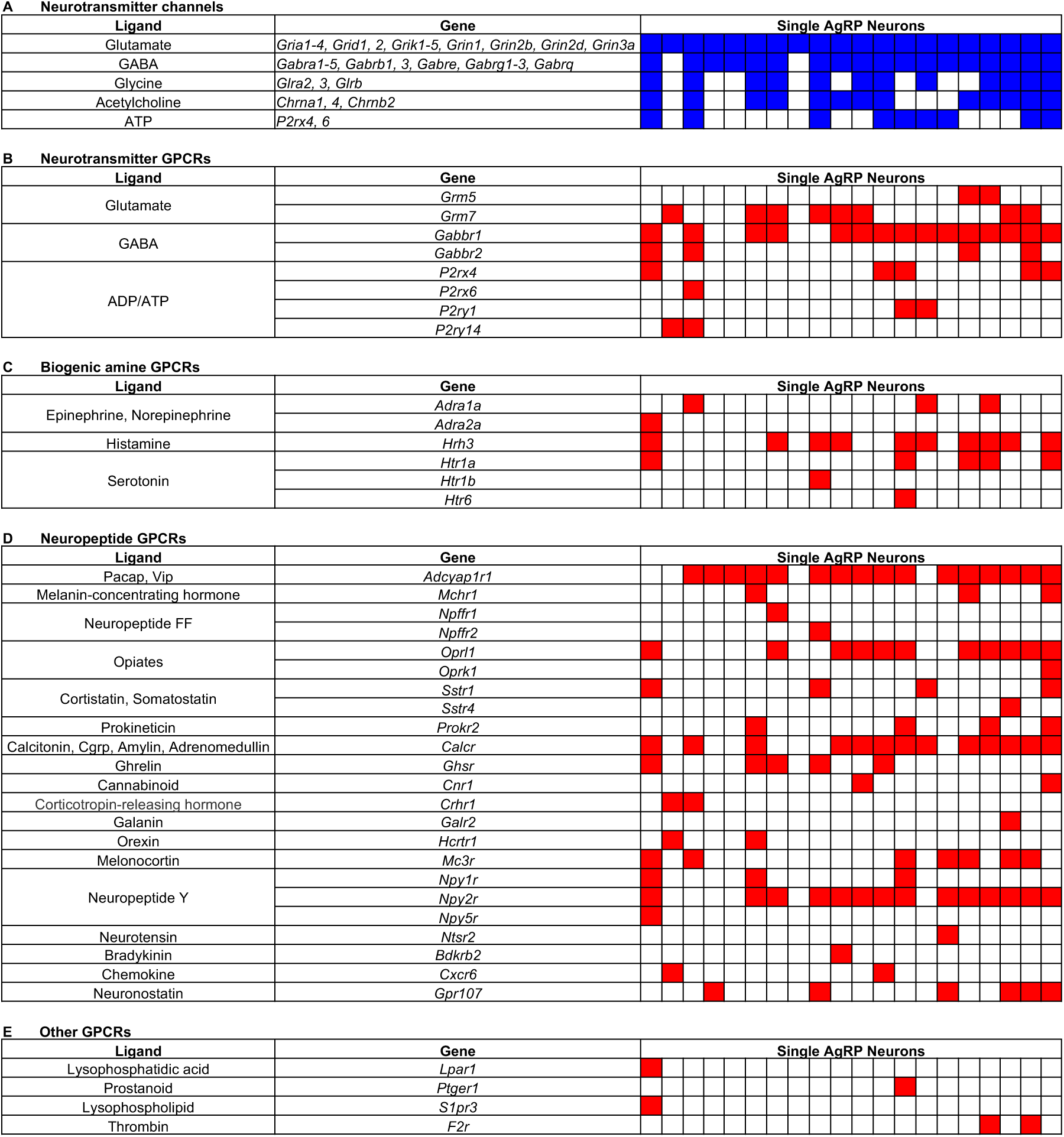
AgRP neurons express multiple receptors for neuromodulators. Transciptome data from individual AgRP neurons indicate that single AgRP neurons can express genes encoding multiple neurotransmitter channels and GPCRs and that those genes are expressed in different combinations in different neurons. Shown here are neurotransmitter channels and GPCRs with known ligands indicated on the left. Colored boxes indicate expression in individual AgRP neurons. Blue boxes show neurotransmitter channels expressed in AgRP neurons (A). GPCRs expressed in AgRP neurons are indicated by red boxes and include GPCRs for neurotransmitters (B), biogenic amines (C), neuropeptides (D), and other ligands (E).

AgRP neurons also exhibited expression of ionotropic (ligand-gated ion channel) receptors for neurotransmitters^37^ (Fig. 5). All expressed ionotropic receptors for glutamate, the major excitatory neurotransmitter in the brain. Many also expressed ionotropic receptors for GABA, the major inhibitory neurotransmitter in the brain. And some also expressed ionotropic receptors for glycine, acetylcholine and/or ATP.

These findings resemble previous RNA-seq data from grouped AgRP neurons^38^. However, the present data go beyond those data by providing information about individual AgRP neurons.

The present data make several important points. First, individual AgRP neurons can respond to multiple different GPCR ligands. On average, each neuron expressed 8 different GPCRs. Most expressed one or more neuropeptide GPCRs. Many also expressed one or more GPCRs for biogenic amines, neurotransmitters, or other ligands.

Second, individual AgRP neurons show considerable heterogeneity in the specific combination of GPCRs they express. Thus, different neurons may respond to different combinations of GPCR ligands.

Third, the prevalence of specific GPCRs varies among neurons. For example, the neuropeptide receptor Adcyap1r1 was found in 80% of neurons while the neuropeptide receptor Galr2 was identified in only 5% (Fig. 5, Fig. S5).

Together, these results suggest that individual AgRP neurons can vary considerably in the combination of GPCRs they express. This variation suggests that they can differ in the incoming signals to which they can respond and, in turn, the signals they send to downstream neurons to modulate appetite.

### Ligands of AgRP neuron receptors define subsets of upstream neurons

The viral tracing experiments identified neurons directly upstream of AgRP neurons in eleven non-olfactory brain areas. The single cell transcriptome experiments revealed AgRP neuron receptors for multiple signaling molecules, including ionotropic neurotransmitter receptors and GPCRs for neurotransmitters, biogenic amines, and neuropeptides. One key question to understanding the neural circuits controlling appetite is whether there are links between the anatomical locations of upstream neurons and the GPCR ligands they use to communicate with AgRP neurons.

To address this question, we employed the Receptor-Assisted Mapping of Upstream Neurons (RAMUN) method we previously used to characterize signaling molecules in neurons upstream of CRH neurons^28^. We infected AgRP neurons with PRVB177, allowed three days for the virus to traverse one synapse, and then costained brain sections for PRV and individual neuropeptides (Fig. 4).

We analyzed the expression of seven neuropeptides in PRV+ neurons directly upstream of AgRP neurons (Fig. 6, Fig. S6). The distribution of these neuropeptides varied widely. For example, Crh was detected exclusively in PRV+ cells within the paraventricular nucleus (PVN), while Pnoc was found in PRV+ cells in six different areas.

**Fig. 6.**
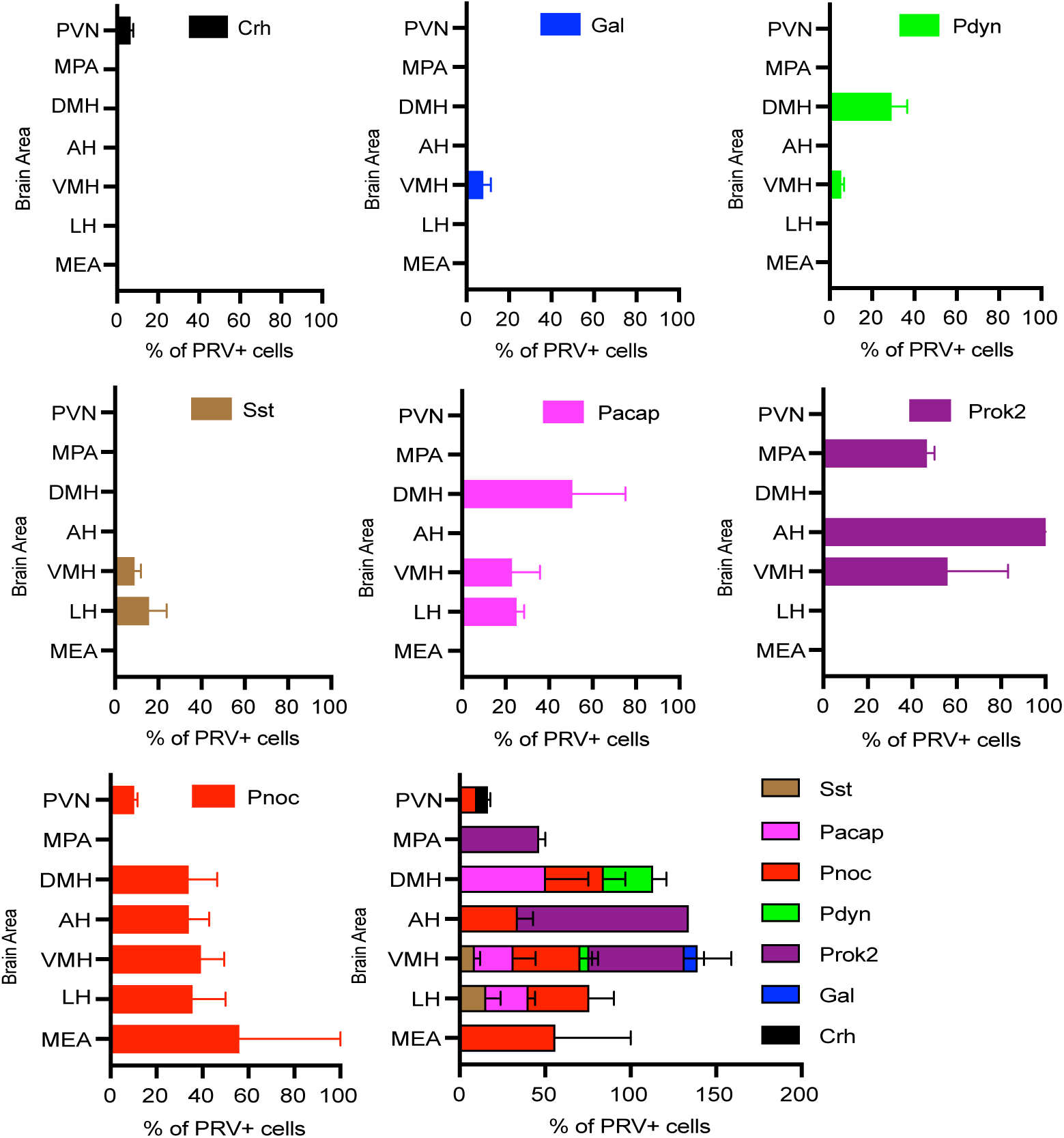
Neuropeptide ligands of AgRP neuron receptors are expressed by upstream neurons in specific areas. Graphs show percentages of PRV+ neurons labeled for specific neuropeptides in different areas on day 3 after infection of AgRP neurons with PRVB177 (n = 3-4). Error bars indicate SEM. See Methods for full names of abbreviated brain areas.

The proportion of PRV+ cells expressing specific neuropeptides also varied. For example, only 6.3 ± 1.6% of PRV+ cells in the PVN expressed Crh, and 7.6 ± 3.8% of PRV+ cells in the ventromedial hypothalamus (VMH) expressed Gal. At the other extreme, every PRV+ neuron in the anterior hypothalamus (AH) expressed Prok2.

In addition, a given neuropeptide was observed in different percentages of PRV+ neurons in different areas. For instance, Pnoc was present in 10.0 ± 1.7% of PRV+ cells in the PVN compared to 55.9 ± 44.1% in the medial amygdala (MEA).

The total percentage of the PRV+ neurons expressing different neuropeptides in DMH, AH, and VMH was over 100%, suggesting that there are upstream neurons expressing more than one of the neuropeptides in these brain areas (Fig. 6). We observed no correlation between the total number of PRV-infected neurons and the percentages of neurons labeled for specific neuropeptides in different areas, indicating that the variations in neuropeptide expression are not simply due to differences in viral infection rates. In addition, there was no apparent correlation between the number of upstream neurons expressing a particular neuropeptide and the proportion of AgRP neurons expressing the corresponding receptor.

Together, these results show that neuropeptide ligands of AgRP neuron GPCRs are differentially expressed in presynaptic neurons in a number of areas, primarily in the hypothalamus. Altogether, seven neuropeptide ligands were seen in six of the eleven areas directly upstream of AgRP neurons. Their distribution within that set of areas varied, but it clearly suggests that the neuropeptide ligands tend to be selectively expressed in upstream neurons in specific brain areas. These results highlight routes of transmission of olfactory and other cues to AgRP neurons and signaling mechanisms employed in those routes to modulate AgRP neurons and their effects on appetite.

## Discussion

The olfactory system has profound effects on appetite and food intake. Here, we sought to gain insight into the mechanisms and neural circuits that underlie these effects. We focused on two subsets of appetite-linked neurons in the hypothalamic arcuate nucleus: AgRP neurons, which stimulate appetite, and POMC neurons, which suppress it.

To examine neural circuits that convey information to AgRP or POMC neurons, they were infected with PRVB177. This virus travels retrogradely across multiple synapses in a time-dependent manner after infecting Cre-expressing neurons.

These studies show that both AgRP and POMC neurons receive information indirectly from the olfactory cortex. The OC contains numerous anatomically distinct areas. Neurons upstream of AgRP or POMC neurons were seen in some of those areas, but not others, suggesting the involvement of only some OC areas in the control of appetite.

The results further indicate that different OC areas communicate with AgRP versus POMC neurons. While one OC area sends information to both, four other OC areas send signals only to AgRP or POMC neurons. Thus, AgRP and POMC neurons are likely to receive different sets of olfactory information.

The respective functions of different OC areas are largely obscure^16^. However, one (AmPir) is linked to physiological fear responses^22^ and another (PLCo) to innate attraction and aversion^39^. It is conceivable that OC areas upstream of AgRP or POMC neurons similarly have distinct effects on appetite, particularly the PLCo, which we saw indirectly upstream of AgRP but not POMC neurons. These considerations emphasize the importance of further exploration to elucidate the roles of different OC areas upstream of AgRP and POMC neurons in appetite regulation.

These analyses extended beyond the main olfactory pathway to consider inputs from the vomeronasal amygdala (VA), which processes olfactory sensory signals about social cues from the vomeronasal organ (VNO)^15,16,24^. Upstream neurons in the two areas of the VA suggest these areas as influencers of both AgRP and POMC neurons. It also opens the possibility that social cues detected by the VNO could have a direct impact on appetite and energy metabolism, integrating social stimuli with fundamental physiological responses

These studies show that AgRP and POMC neurons receive direct input from numerous non-olfactory brain areas, primarily in the hypothalamus. These neurons have the potential to relay information from the olfactory cortex to the appetite neurons. Most or all projection neurons in the olfactory cortex are excitatory glutamatergic neurons^22^. However, they could activate either excitatory or inhibitory neurons presynaptic to AgRP or POMC neurons and thereby cause either activation or suppression of the appetite neurons.

In exploring the molecular basis of these connections, we discovered a rich diversity of GPCRs in AgRP neurons. These receptors, which include those for neurotransmitters like glutamate and GABA, as well as for various neuropeptides, illustrate the complexity of input integration at the neuronal level. The differential expression of these GPCRs among AgRP neurons suggests a mechanism where individual AgRP neurons may be selectively responsive to a limited spectrum of those signaling molecules, potentially directing varied physiological responses.

The importance of the AgRP neuron GPCRs and GPCR ligands revealed in these experiments is supported by genetic studies that link these molecules to obesity and energy homeostasis in humans or mice^40^. These include human obesity-associated polymorphisms in three receptors: CRH1^41^, MCHR1^42–44^, and MC3R^45–48^. In mice, they include three receptors whose deletion increases body weight and adiposity (Mc3r^49,50^, Npy1r^51^, and Npy5r^52^) and two that have the opposite effect (Ghsr^53,54^ and Cnr1^55^). The ligands of identified receptors also have effects when modulated, with overexpression of Crh increasing body weight^56^ and deletion of Adcyap1 decreasing it^57^. These associations between AgRP neuron receptors and their ligands and genes that alter energy homeostasis emphasize the potential importance of these molecules as pharmaceutical targets.

By defining the specific ligands of AgRP neuron receptors and the distinct subsets of upstream neurons that express these ligands, these studies uncovered molecular genetic tools to further dissect the roles of these neurons in controlling appetite. This foundational knowledge sets the stage for future studies aimed at understanding how different neural circuits integrate multiple forms of input, including olfactory signals, to regulate appetite. It also provides a rich complement of receptors and ligands that offer potential therapeutic targets for disorders of appetite and energy homeostasis.

## Methods

### Animals

C57BL/6J wildtype, AgRP-IRES-Cre^33^ (Jax stock no: 012899), Ai6^34^ (Jax stock#: 007906), Pomc-eGFP^58^ (Jax stock#: 009593), and POMC-Cre^59^ (Jax stock#: 005965) mice were purchased from the Jackson Laboratory. All procedures involving mouse handling were approved by the Fred Hutchinson Cancer Center Institutional Animal Care and Use Committee.

### Viral vectors

PRVB177 was constructed as previously described^22^. Briefly, a CMV promoter, a flexstop-flanked sequence encoding a PRV thymidine kinase (TK) fused with a hemagglutinin (HA) epitope tag, and a SV40 polyadenylation signal, were cloned into PRV TK-BaBlu, a TK-deleted PRV Bartha strain between its gG locus sequences matching 5′ and 3′ to the lacZ sequence. Recombinant virus clones were selected and confirmed as described previously^22^.

Recombinant PRVs were propagated in PK15 cells (ATCC) using a multiplicity of infection (M.O.I.) = 0.1∼0.01. Three days after infection, cells were harvested by scraping. Material was frozen using liquid nitrogen and then quickly thawed in a 37°C water bath 3 times. Cell debris was then removed by centrifugation twice at 1,000xg for 5 minutes. The titer of supernatant was determined using standard plaque assays on PK15 cells, with titers expressed in plaque-forming units (PFU).

### Stereotaxic injection

Stereotaxic injection was performed as previously described^22^. Virus suspension (1–2 × 10^9^ PFU) was injected into the brains of mice aged 2-6 months using a 2 μl syringe at 100 nl per minute. Targeted brain areas were referenced based on a stereotaxic atlas^60^ using a Stereotaxic Alignment System (David Kopf Instruments). Mice were treated with an inhalation anesthesia of 2.5% Isoflurane during injection. Animals were singly housed with regular 12 h dark/light cycles in the presence of food and water ad libitum after recovery.

### Staining for neuropeptide mRNA and PRVB177 (HA)

Experiments were performed as described previously^22^. For perfusion, 4% paraformaldehyde (PFA) was used to perfuse animals transcardially. Brains were dissected out and then soaked in 4% PFA overnight. After soaking in 30% sucrose for 48 hours, the brains were frozen in OCT (Sakura) and stored at −80 °C before sectioning. For fresh frozen tissues, animals were decapitated immediately after cervical dislocation. Fresh brains were dissected out, snap frozen in isopentane mixed with dry ice, and kept at −80 °C. Brains were sectioned into 20 μm coronal sections using a cryostat.

For detection of PRV positive neurons, brain sections were incubated with biotinylated mouse anti-HA antibodies (BioLegend, #901505, 1:300) at 4°C overnight or at 37°C for an hour. Sections were then incubated with 0.5 μg/ml 4’, 6-diamidino-2-phenylindole (DAPI, Sigma) and Alexa488 Streptavidin (Thermo Fisher, 1:1000) at room temperature for 1 hour followed by coverslipping with Fluoromount-G (Southern Biotech). For double staining of PRV-infected neurons expressing neuropeptides, coding regions of neuropeptide genes were amplified from mouse brain cDNA using PCR and cloned into the pCR4 TOPO vector (Thermo Fisher). Digoxigenin (DIG)-labeled cRNA probes (riboprobes) were prepared using the DIG RNA Labeling Mix (Roche). Twenty μm coronal cryostat sections of mouse brains frozen in OCT were hybridized to DIG-labeled cRNA probes at 56°C for 13–16 hours. After hybridization, brain sections were washed in 5xSSC and 0.2×SSC at 63°C for 30 minutes each consecutively. Sections were incubated with POD-conjugated anti-DIG antibodies (Roche, #11207733910, 1:2000) and biotinylated anti-HA antibodies (BioLegend, #901505, 1:300) at 4°C for overnight or at 37°C for an hour and treated using the TSA-plus Cy3 kit (Perkin Elmer) according to manufacturer’s instruction. Sections were then incubated with 0.5 μg/ml DAPI and Alexa488 Streptavidin (Thermo Fisher, 1:1000) at room temperature for 1 hour. Slides were coverslipped with Fluoromount-G.

### Dissociation of AgRP neurons and RNA-sequencing

AgRP-IRES-Cre mice^33^ were crossed with Ai6 mice^34^, which have Cre-dependent expression of eGFP, to generate AgRP-Cre:Ai6 mice. Arcuate nuclei of AgRP-Cre:Ai6 mice were then dissected out as previously described^36^ to obtain AgRP+ cells. GFP-labelled single cells were manually isolated from AgRP-Cre:Ai6 mice using a micropipette and visualizing cells under a fluorescent microscope. Single cells were lysed, and cDNA libraries were generated similar to our previously published protocols^28,35,36^. A total of 20 cells with good quality cDNAs were analyzed to confirm the expression of AgRP and NPY. The following intron-spanning primers were used: AgRP-5’, CAACTGCAGACCGAGCAGAAG, AgRP-3’-GCAGCAAGGTACCTGCTGTC, NPY-5’, GGCACCCAGAGCAGAGCAC, and NPY-3’, CCAGAATGCCCAAACACACG. cDNAs that confirmed expression of markers were processed for Illumina sequencing similar to previously published methods^36^. Sequenced reads were initially mapped to the mouse genome to generate reads per million (RPM) data. Later, Fragments Per Kilobase of exon per Million mapped fragments (FPKM) data were also used with standard methods (Tophat and Cufflinks)^61,62^. The list of GPCRs was obtained from the International Union of Basic and Clinical Pharmacology (IUPHAR)/ British Pharmacological Society (BPS) website (http://www.guidetopharmacology.org).

### Cell counting

Stained brain sections were imaged with AxioImager.Z1 microscope. Brain structures were identified based on a mouse brain atlas^60^. Numbers of PRV-infected cells in brain areas of every fifth section were counted. To acquire approximate total number of cells in each brain area of an animal, brain areas were judged to contain upstream neurons if they contained ≥ 2 labeled neurons in ≥ 50% of animals. All data are shown as the mean±SEM. Neuropeptide counts were confirmed with TissueFax in Fred Hutch Cancer Center Shared Resources.

## Acknowledgments

We thank R. Basom at the Fred Hutchinson Cancer Center (FHCC) Genomics and Bioinformatics Shared Resource for assistance with analyzing RNA-seq data. We are also grateful to members of the Buck Laboratory for helpful discussions and comments.

## Author contributions

D.K., N.K.H., and L.B.B. conceived the project. D.K. conducted and analyzed all the viral tracing experiments and investigation of neuropeptides in upstream neurons and determined ligands of AgRP neuron receptors. N.K.H. isolated single cells, prepared cDNA libraries and conducted the transcriptome analyses of single AgRP neurons and identified receptors for neuromodulators. C-Y.L and A.H. injected animals with virus, X.Y. stained brain sections, X.Y. and M.E. analyzed sections with TissueFax, and D.K., N.K.H., and L.B.B. wrote the manuscript.

## Data and materials availability

All data needed to evaluate the conclusions in the paper are present in the paper. Additional data and materials related to this paper may be requested from the authors. Raw sequencing data related to this study will be archived in the National Center for Biotechnology Gene Expression Omnibus (GEO) database prior to formal publication.

---

**Fig. S1.**
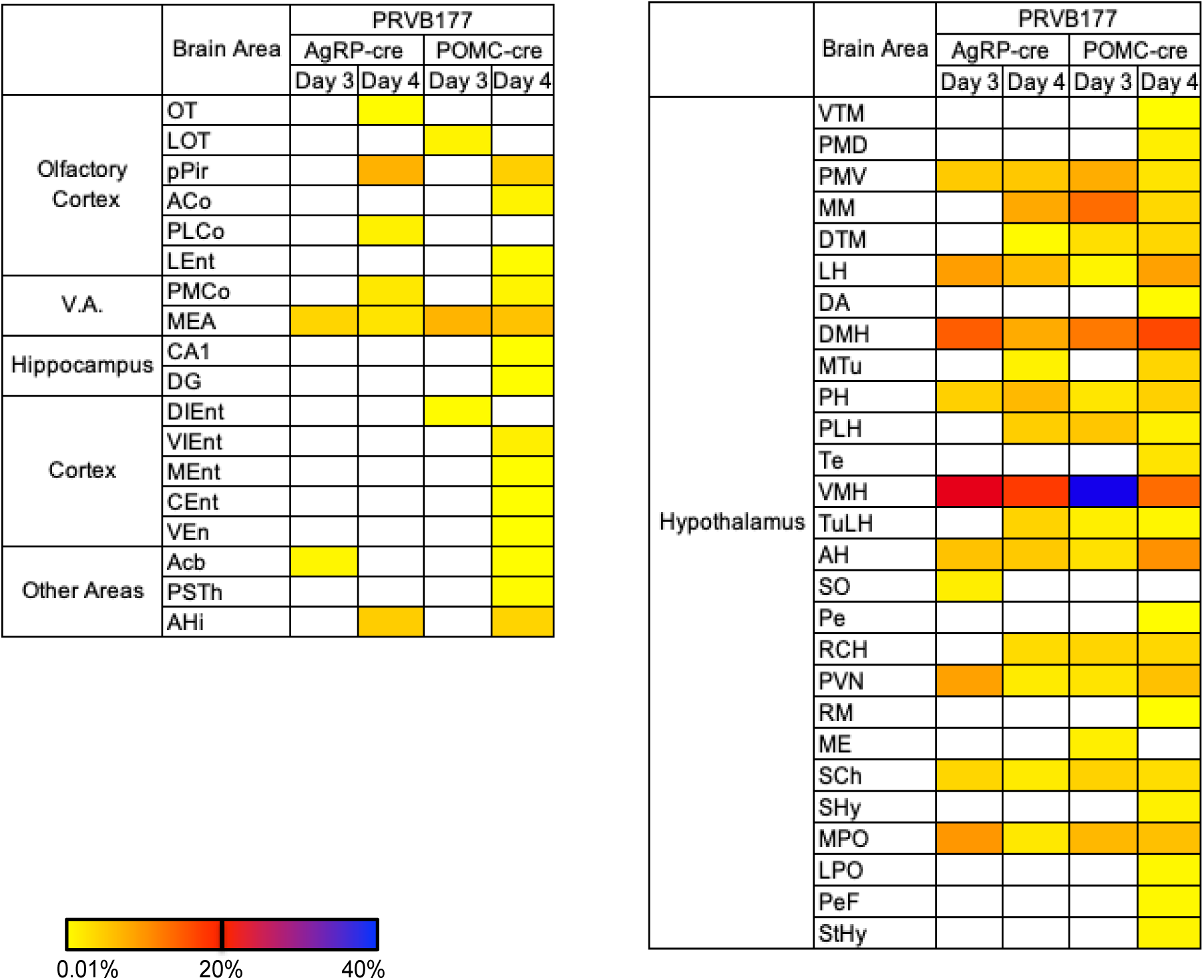
Heat map of brain areas with neurons upstream of AgRP or POMC neurons. Colored boxes indicate approximate percentages of all non-ARC PRV+ neurons in individual brain areas, denoted by color intensity (white indicates none) on day 3 or 4 after ARC injection of AgRP-Cre or POMC-Cre mice with PRVB177. Sample sizes as in Fig. 2. Areas with PRV+ neurons on day 3 are directly upstream of AgRP or POMC neurons. Areas with PRV+ neurons on day 4 but not day 3 are indirectly upstream of AgRP or POMC neurons. See Methods for full names of abbreviated brain areas.

**Fig. S2.**
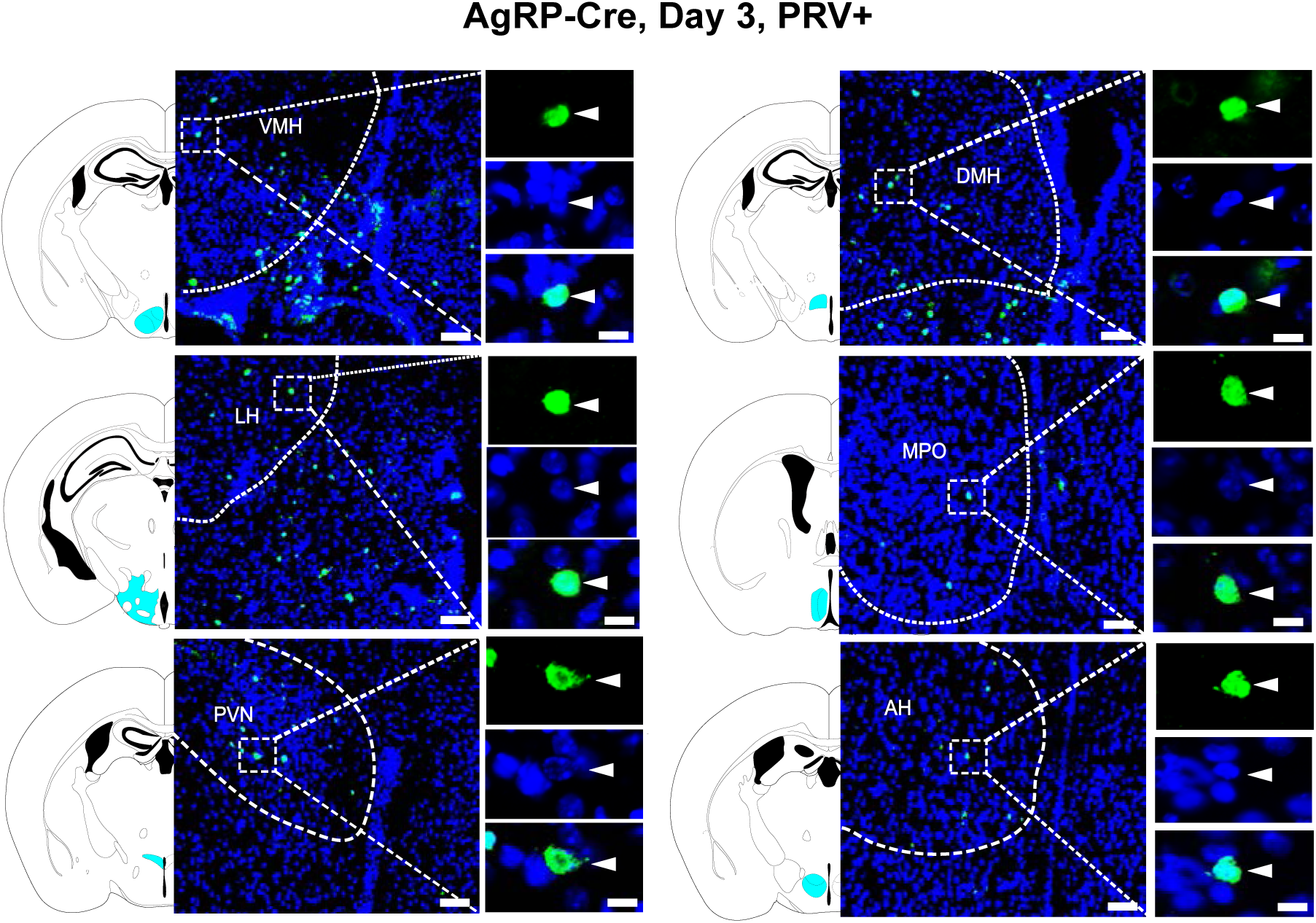
Non-olfactory areas with PRV+ neurons directly upstream of AgRP neurons. Images of cells immunostained for PRV (HA) (green) in different non-olfactory areas on day 3 post-injection of AgRP-Cre mice. DAPI counterstain, blue. Corresponding areas on diagrams are labeled with cyan. Dotted lines indicate locations of brain areas. Boxed areas are shown at higher magnification at right (top, PRV; middle, DAPI; bottom, merged). Scale bars, 100 μm (middle) and 20 μm (right).

**Fig. S3.**
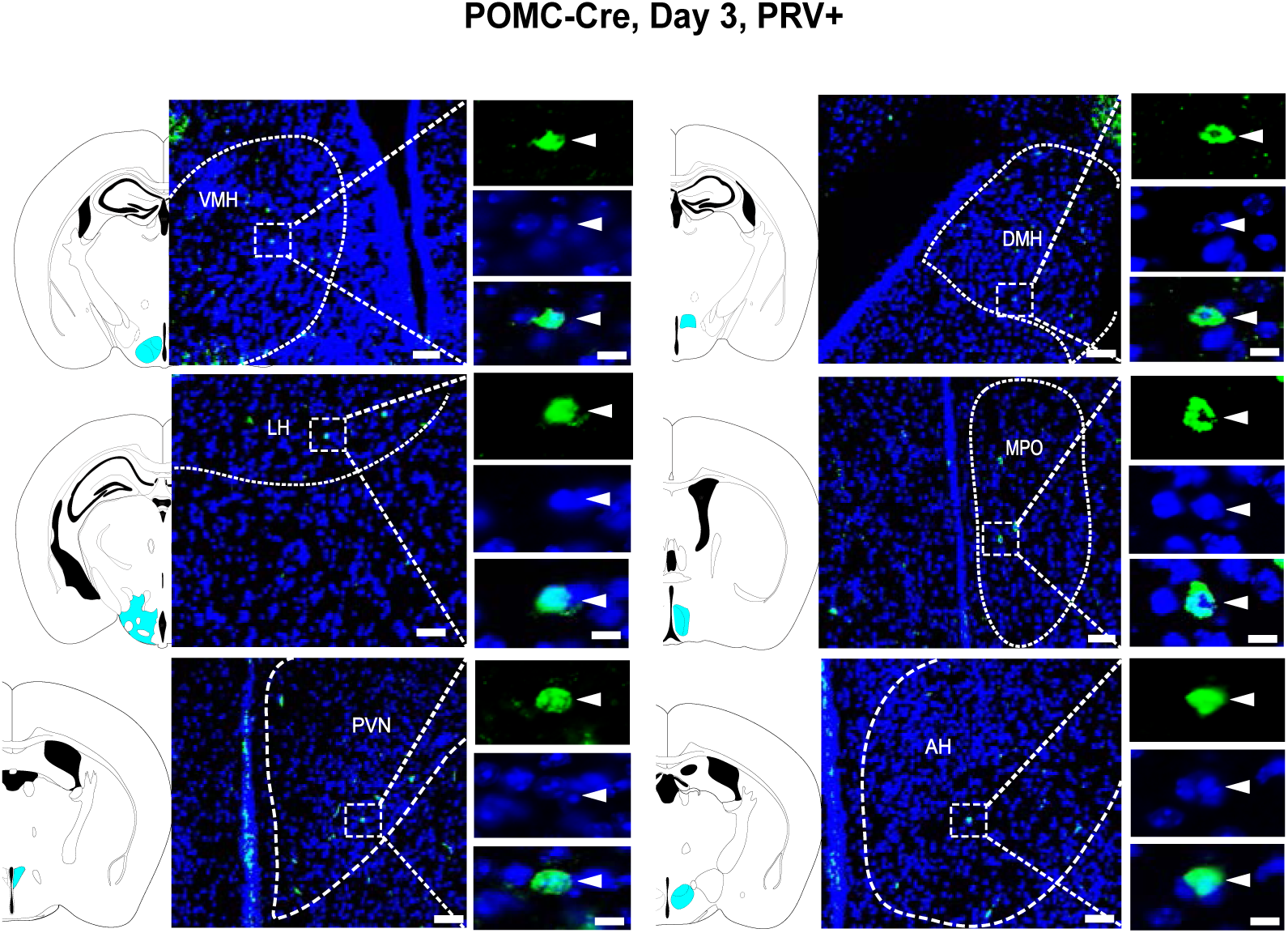
Non-olfactory areas with PRV+ neurons directly upstream of POMC neurons. Images of cells immunostained for PRV (HA) (green) in different non-olfactory areas on day 3 post-injection of POMC-Cre mice. DAPI counterstain, blue. Corresponding areas on diagrams are labeled with cyan. Dotted lines indicate locations of brain areas. Boxed areas are shown at higher magnification at right (top, PRV; middle, DAPI; bottom, merged). Scale bars, 100 μm (middle) and 20 μm (right).

**Fig. S4.**
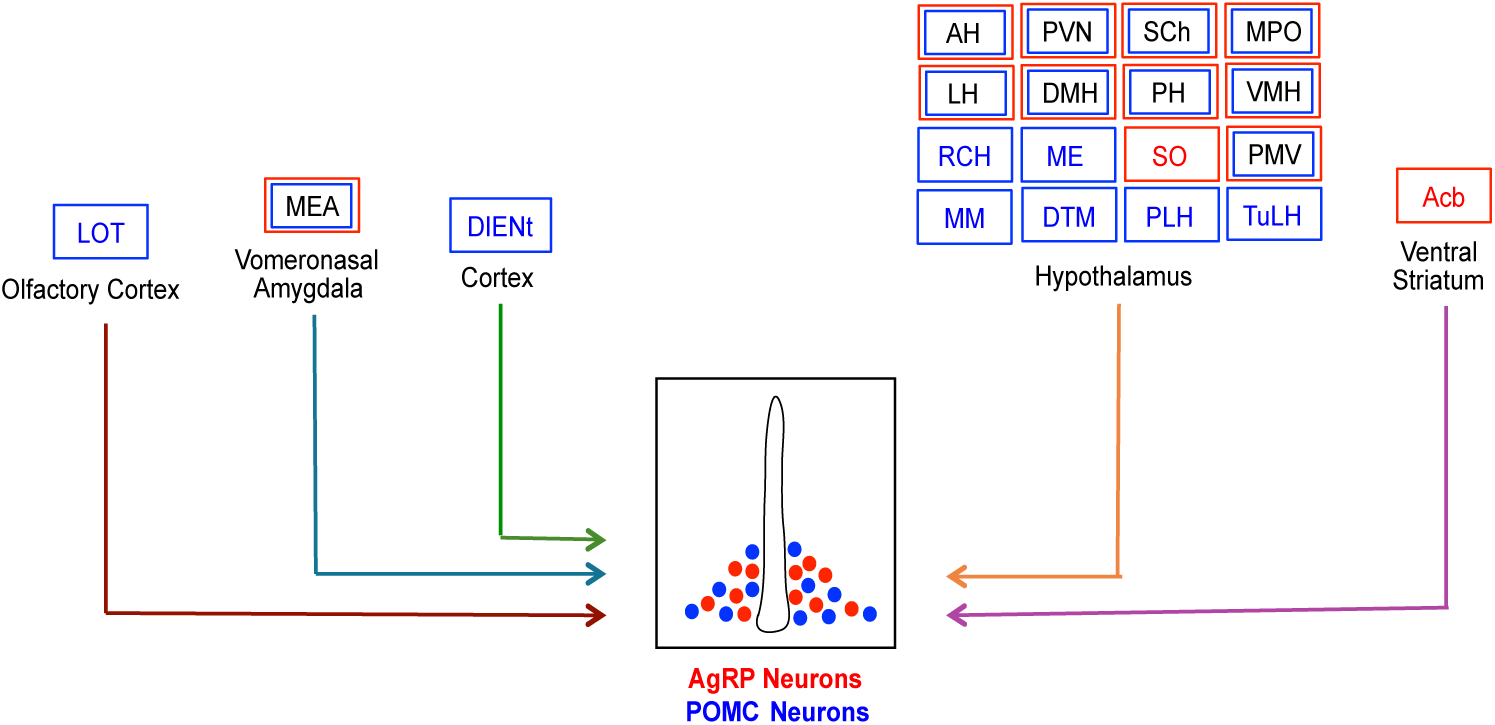
Anatomical map of direct inputs to AgRP and POMC neurons. Schematic shows brain areas containing neurons directly upstream of AgRP or POMC neurons, illustrated in red and blue, respectively. Twelve areas (including 11 non-olfactory) contain neurons upstream of AgRP neurons while eighteen (including 16 non-olfactory) areas contain neurons upstream of POMC neurons. Ten areas (including 9 non-olfactory) contain neurons directly upstream of both AgRP and POMC neurons.

**Fig. S5.**
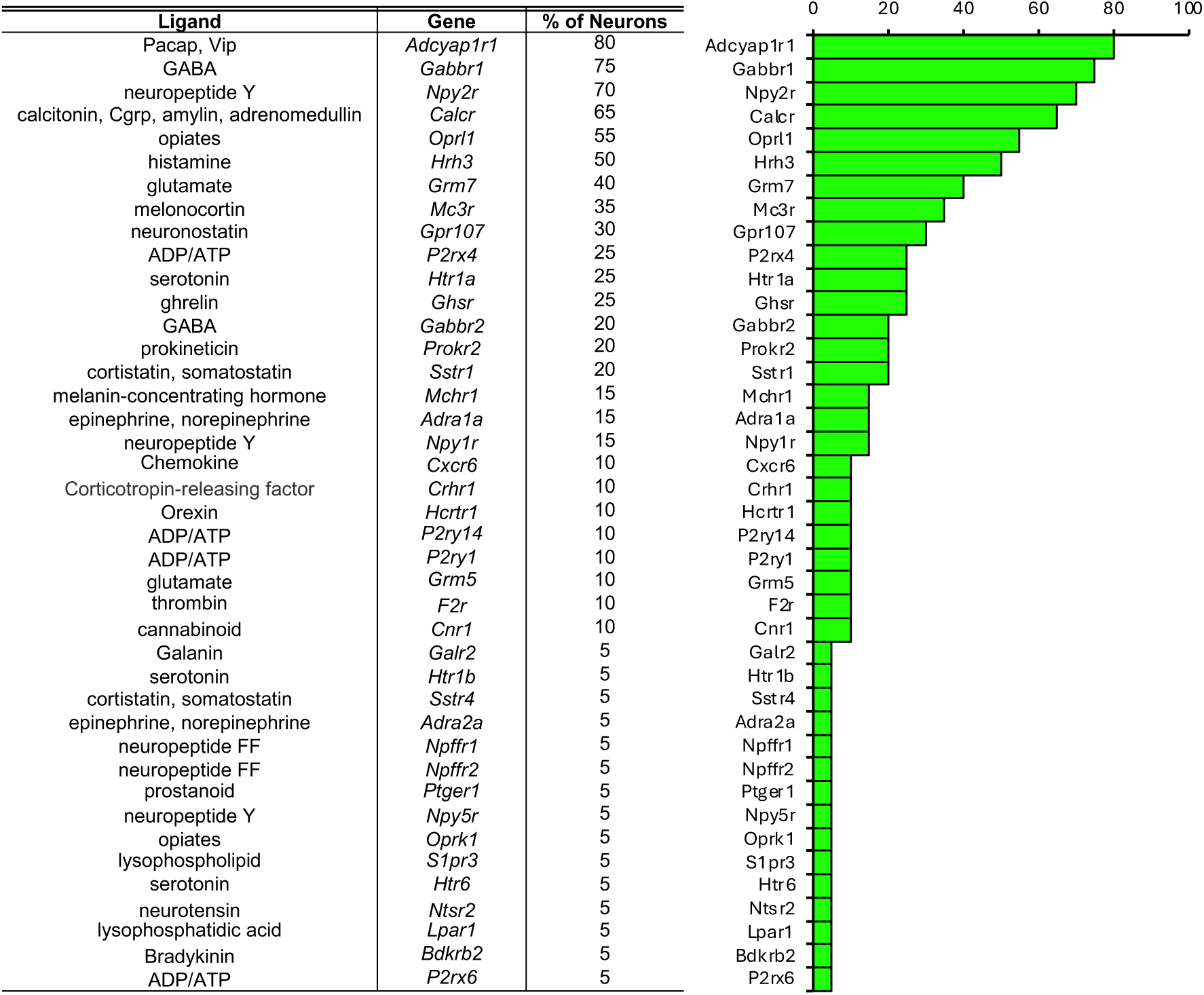
GPCR expression in scRNA-seq data. This chart displays the percentages of AgRP neurons in the scRNA-seq dataset that express genes encoding GPCRs with known ligands, as indicated. The percentages are visually represented as green bars on the right.

**Fig. S6.**
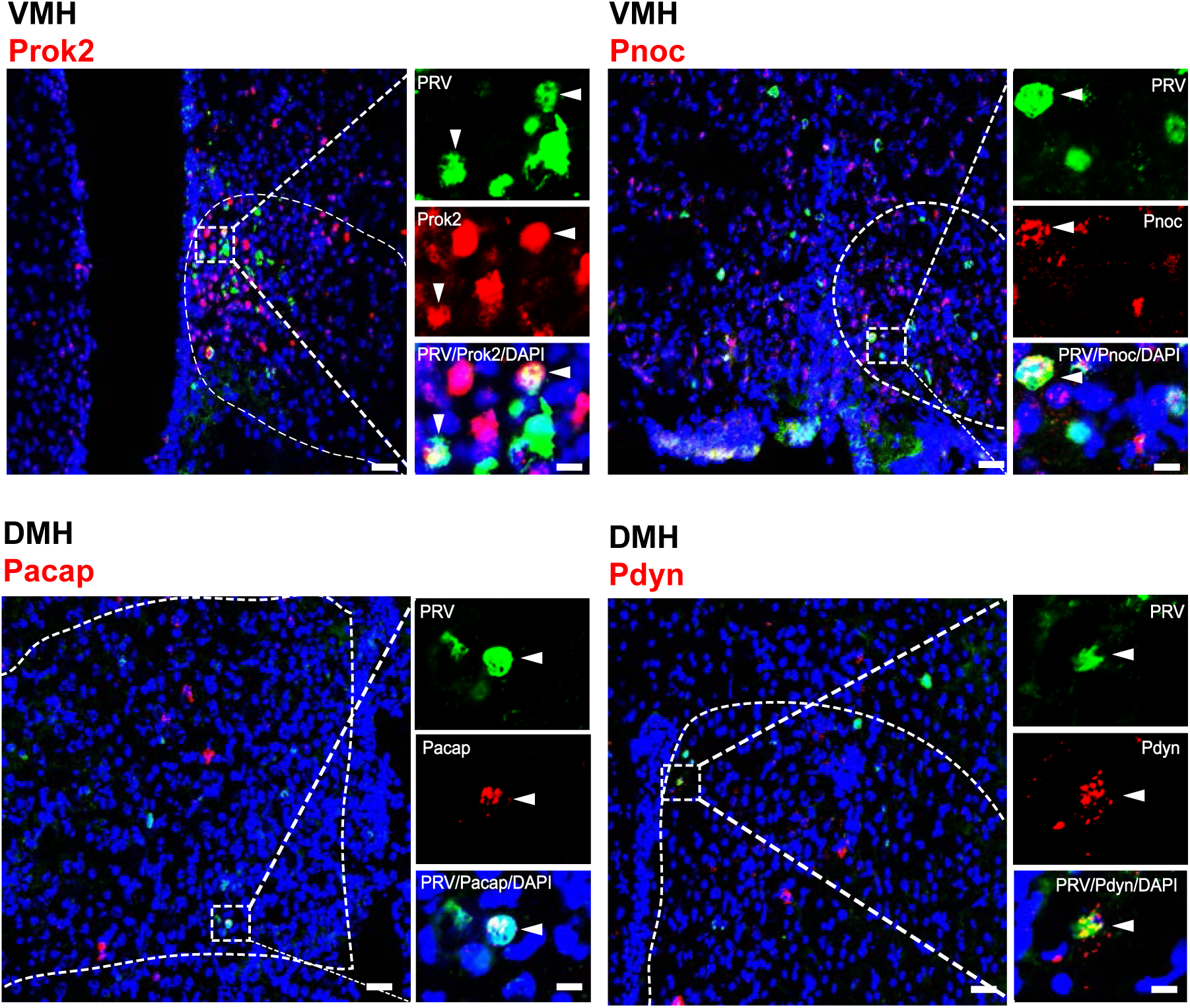
Expression of individual neuropeptides in single PRV+ cells in AgRP-Cre mice. Photographs show expression of individual neuropeptides (Prok2, Pnoc, Pacap, or Pdyn) in single PRV+ neurons in VMH or DMH. PRV+, green; neuropeptide+, red; DAPI+, blue. Brain areas are indicated by dotted lines. Boxed areas are shown at higher magnification at right. Scale bars, 100 μm (left) and 20 μm (right).

## Abbreviations for brain areas

Modified based on Franklin and Paxinos^60^.

Acb: accumbens nucleus
ACo: anterior cortical amygdaloid area
AH: anterior hypothalamic area
AHi: amygdalohippocampal area
AmPir: amygdalo-piriform transition area
AON: anterior olfactory nucleus
aPir: piriform cortex, anterior part
ARC: arcuate hypothalamic nucleus
CA1: field CA1 of the hippocampus
CEnt: caudomedial entorhinal cortex
Coxe: cortex-amygdala transition zone
DA: dorsal hypothalamic area
DG: dentate gyrus
DIEnt: dorsal intermediate entorhinal cortex
DMH: dorsomedial hypothalamic nucleus
DTM: dorsal tuberomammillary nucleus
LEnt: lateral entorhinal cortex
LH: lateral hypothalamic area
LOT: nucleus of the lateral olfactory tract
LPO: lateral preoptic area
ME: median eminence
MEA: medial amygdala
MEnt: medial entorhinal cortex
MM: medial mammillary nucleus, medial part
MPA: medial preoptic area
MPO: medial preoptic nucleus
MTu: medial tuberal nucleus
OC: olfactory cortex
OT: olfactory tubercle
Pe: periventricular hypothalamic nucleus
PeF: perifornical nucleus
PH: posterior hypothalamic nucleus
PLCo: posterolateral cortical amygdaloid area
PLH: peduncular part of lateral hypothalamus
PMCo: posteromedial cortical amygdala
PMD: premammillary nucleus, dorsal part
PMV: premammillary nucleus, ventral part
pPir: piriform cortex, posterior part
PSTh: posterior subthalamic nucleus
PVN: paraventricular nucleus of the hypothalamus
RCH: retrochiasmatic area
RM: retromammillary nucleus
SCh: suprachiasmatic nucleus
SHy: septohypothalamic nucleus
SO: supraoptic nucleus
StHy: striohypothalamic nucleus
Te: terete hypothalamic nucleus
TT: tenia tecta
TuLH: tuberal region of lateral hypothalamus
VA: vomeronasal amygdala
VEn: ventral endopiriform claustrum
VIEnt: ventral intermediate entorhinal cortex
VMH: ventromedial hypothalamic nucleus
VTM: ventral tuberomammillary nucleus

## References

1. Andermann, M.L. & Lowell, B.B. Toward a Wiring Diagram Understanding of Appetite Control. Neuron 95, 757–778 (2017).

2. Jais, A. & Bruning, J.C. Arcuate Nucleus-Dependent Regulation of Metabolism-Pathways to Obesity and Diabetes Mellitus. Endocr Rev 43, 314–328 (2022).

3. Hahn, T.M., Breininger, J.F., Baskin, D.G. & Schwartz, M.W. Coexpression of Agrp and NPY in fasting-activated hypothalamic neurons. Nat Neurosci 1, 271–272 (1998).

4. Schwartz, M.W., Erickson, J.C., Baskin, D.G. & Palmiter, R.D. Effect of fasting and leptin deficiency on hypothalamic neuropeptide Y gene transcription in vivo revealed by expression of a lacZ reporter gene. Endocrinology 139, 2629–2635 (1998).

5. Wu, Q., et al. The temporal pattern of cfos activation in hypothalamic, cortical, and brainstem nuclei in response to fasting and refeeding in male mice. Endocrinology 155, 840–853 (2014).

6. Aponte, Y., Atasoy, D. & Sternson, S.M. AGRP neurons are sufficient to orchestrate feeding behavior rapidly and without training. Nat Neurosci 14, 351–355 (2011).

7. Krashes, M.J., et al. Rapid, reversible activation of AgRP neurons drives feeding behavior in mice. J Clin Invest 121, 1424–1428 (2011).

8. Dietrich, M.O., Zimmer, M.R., Bober, J. & Horvath, T.L. Hypothalamic Agrp Neurons Drive Stereotypic Behaviors beyond Feeding. Cell 160, 1222–1232 (2015).

9. Cavalcanti-de-Albuquerque, J.P., Bober, J., Zimmer, M.R. & Dietrich, M.O. Regulation of substrate utilization and adiposity by Agrp neurons. Nat Commun 10, 311 (2019).

10. Wei, Q., et al. Uneven balance of power between hypothalamic peptidergic neurons in the control of feeding. Proc Natl Acad Sci U S A 115, E9489–e9498 (2018).

11. Betley, J.N., et al. Neurons for hunger and thirst transmit a negative-valence teaching signal. Nature 521, 180–185 (2015).

12. Chen, Y., Lin, Y.C., Kuo, T.W. & Knight, Z.A. Sensory detection of food rapidly modulates arcuate feeding circuits. Cell 160, 829–841 (2015).

13. Mandelblat-Cerf, Y., et al. Arcuate hypothalamic AgRP and putative POMC neurons show opposite changes in spiking across multiple timescales. Elife 4(2015).

14. Su, Z., Alhadeff, A.L. & Betley, J.N. Nutritive, Post-ingestive Signals Are the Primary Regulators of AgRP Neuron Activity. Cell Rep 21, 2724–2736 (2017).

15. L. Buck, K.S., C. Zuker. Smell and Taste: the Chemical Senses, (McGraw-Hill, 2021).

16. Luo, L. Principles of Neurobiology, (CRC Press, 2021).

17. Haberly, L.B. Parallel-distributed processing in olfactory cortex: new insights from morphological and physiological analysis of neuronal circuitry. Chem Senses 26, 551–576 (2001).

18. Pashkovski, S.L., et al. Structure and flexibility in cortical representations of odour space. Nature 583, 253–258 (2020).

19. Fulton, K.A., Zimmerman, D., Samuel, A., Vogt, K. & Datta, S.R. Common principles for odour coding across vertebrates and invertebrates. Nat Rev Neurosci 25, 453–472 (2024).

20. Krashes, M.J., et al. An excitatory paraventricular nucleus to AgRP neuron circuit that drives hunger. Nature 507, 238–242 (2014).

21. Wang, D., et al. Whole-brain mapping of the direct inputs and axonal projections of POMC and AgRP neurons. Front Neuroanat 9, 40 (2015).

22. Kondoh, K., et al. A specific area of olfactory cortex involved in stress hormone responses to predator odours. Nature 532, 103–106 (2016).

23. Buck, L.B., and Bargmann, C. Smell and Taste: the Chemical Senses. in Principles of Neuroscience (ed. Kandel, E., Schwartz, J., Jessell, T., Siegelbaum, S., Hudspeth, A.J.) 712-742 (McGraw-Hill, New York, 2012).

24. Mohrhardt, J., Nagel, M., Fleck, D., Ben-Shaul, Y. & Spehr, M. Signal Detection and Coding in the Accessory Olfactory System. Chem Senses 43, 667–695 (2018).

25. Deem, J.D., Faber, C.L. & Morton, G.J. AgRP neurons: Regulators of feeding, energy expenditure, and behavior. FEBS J 289, 2362–2381 (2022).

26. Schwartz, M.W. Central nervous system regulation of food intake. Obesity (Silver Spring) 14 Suppl 1, 1S–8S (2006).

27. Bains, J.S., Wamsteeker Cusulin, J.I. & Inoue, W. Stress-related synaptic plasticity in the hypothalamus. Nat Rev Neurosci 16, 377–388 (2015).

28. Lee, E.J., et al. A psychological stressor conveyed by appetite-linked neurons. Sci Adv 6, eaay5366 (2020).

29. Ulrich-Lai, Y.M. & Herman, J.P. Neural regulation of endocrine and autonomic stress responses. Nat Rev Neurosci 10, 397–409 (2009).

30. Lee, E.J., et al. Odor blocking of stress hormone responses. Sci Rep 12, 8773 (2022).

31. Zimmerman, C.A., Leib, D.E. & Knight, Z.A. Neural circuits underlying thirst and fluid homeostasis. Nat Rev Neurosci 18, 459–469 (2017).

32. Klawonn, A.M. & Malenka, R.C. Nucleus Accumbens Modulation in Reward and Aversion. Cold Spring Harb Symp Quant Biol 83, 119–129 (2018).

33. Tong, Q., Ye, C.-P., Jones, J.E., Elmquist, J.K. & Lowell, B.B. Synaptic release of GABA by AgRP neurons is required for normal regulation of energy balance. Nat Neurosci 11, 998–1000 (2008).

34. Madisen, L., et al. A robust and high-throughput Cre reporting and characterization system for the whole mouse brain. Nat Neurosci 13, 133–140 (2010).

35. Hanchate, N.K., et al. Single-cell transcriptomics reveals receptor transformations during olfactory neurogenesis. Science 350, 1251–1255 (2015).

36. Hanchate, N.K., et al. Connect-seq to superimpose molecular on anatomical neural circuit maps. Proc Natl Acad Sci U S A 117, 4375–4384 (2020).

37. Southan, C., et al. The IUPHAR/BPS Guide to PHARMACOLOGY in 2016: towards curated quantitative interactions between 1300 protein targets and 6000 ligands. Nucleic Acids Res 44, D1054–D1068 (2016).

38. Henry, F.E., Sugino, K., Tozer, A., Branco, T. & Sternson, S.M. Cell type-specific transcriptomics of hypothalamic energy-sensing neuron responses to weight-loss. Elife 4(2015).

39. Root, C.M., Denny, C.A., Hen, R. & Axel, R. The participation of cortical amygdala in innate, odour-driven behaviour. Nature 515, 269–273 (2014).

40. Rankinen, T., et al. The human obesity gene map: the 2005 update. Obesity (Silver Spring) 14, 529–644 (2006).

41. Challis, B.G., et al. Genetic variation in the corticotrophin-releasing factor receptors: identification of single-nucleotide polymorphisms and association studies with obesity in UK Caucasians. Int J Obes Relat Metab Disord 28, 442–446 (2004).

42. Gibson, W.T., et al. Melanin-concentrating hormone receptor mutations and human obesity: functional analysis. Obes Res 12, 743–749 (2004).

43. Bell, C.G., et al. Association of melanin-concentrating hormone receptor 1 5’ polymorphism with early-onset extreme obesity. Diabetes 54, 3049–3055 (2005).

44. Wermter, A.K., et al. Mutation analysis of the MCHR1 gene in human obesity. Eur J Endocrinol 152, 851–862 (2005).

45. Lee, Y.S., Poh, L.K. & Loke, K.Y. A novel melanocortin 3 receptor gene (MC3R) mutation associated with severe obesity. J Clin Endocrinol Metab 87, 1423–1426 (2002).

46. Rached, M., Buronfosse, A., Begeot, M. & Penhoat, A. Inactivation and intracellular retention of the human I183N mutated melanocortin 3 receptor associated with obesity. Biochim Biophys Acta 1689, 229–234 (2004).

47. Tao, Y.X. & Segaloff, D.L. Functional characterization of melanocortin-3 receptor variants identify a loss-of-function mutation involving an amino acid critical for G protein-coupled receptor activation. J Clin Endocrinol Metab 89, 3936–3942 (2004).

48. Boucher, N., et al. A +2138InsCAGACC polymorphism of the melanocortin receptor 3 gene is associated in human with fat level and partitioning in interaction with body corpulence. Mol Med 8, 158–165 (2002).

49. Butler, A.A., et al. A unique metabolic syndrome causes obesity in the melanocortin-3 receptor-deficient mouse. Endocrinology 141, 3518–3521 (2000).

50. Chen, A.S., et al. Inactivation of the mouse melanocortin-3 receptor results in increased fat mass and reduced lean body mass. Nat Genet 26, 97–102 (2000).

51. Kushi, A., et al. Obesity and mild hyperinsulinemia found in neuropeptide Y-Y1 receptor-deficient mice. Proc Natl Acad Sci U S A 95, 15659–15664 (1998).

52. Marsh, D.J., Hollopeter, G., Kafer, K.E. & Palmiter, R.D. Role of the Y5 neuropeptide Y receptor in feeding and obesity. Nat Med 4, 718–721 (1998).

53. Sun, Y., Wang, P., Zheng, H. & Smith, R.G. Ghrelin stimulation of growth hormone release and appetite is mediated through the growth hormone secretagogue receptor. Proc Natl Acad Sci U S A 101, 4679–4684 (2004).

54. Lall, S., et al. Physiological studies of transgenic mice overexpressing growth hormone (GH) secretagogue receptor 1A in GH-releasing hormone neurons. Endocrinology 145, 1602–1611 (2004).

55. Ravinet Trillou, C., Delgorge, C., Menet, C., Arnone, M. & Soubrie, P. CB1 cannabinoid receptor knockout in mice leads to leanness, resistance to diet-induced obesity and enhanced leptin sensitivity. Int J Obes Relat Metab Disord 28, 640–648 (2004).

56. Stenzel-Poore, M.P., Cameron, V.A., Vaughan, J., Sawchenko, P.E. & Vale, W. Development of Cushing’s syndrome in corticotropin-releasing factor transgenic mice. Endocrinology 130, 3378–3386 (1992).

57. Gray, S.L., Cummings, K.J., Jirik, F.R. & Sherwood, N.M. Targeted disruption of the pituitary adenylate cyclase-activating polypeptide gene results in early postnatal death associated with dysfunction of lipid and carbohydrate metabolism. Mol Endocrinol 15, 1739–1747 (2001).

58. Cowley, M.A., et al. Leptin activates anorexigenic POMC neurons through a neural network in the arcuate nucleus. Nature 411, 480–484 (2001).

59. Balthasar, N., et al. Leptin Receptor Signaling in POMC Neurons Is Required for Normal Body Weight Homeostasis. Neuron 42, 983–991 (2004).

60. Franklin, K. & Paxinos, G. The Mouse Brain in Stereotaxic Coordinates, (Elsevier Inc., 2008).

61. Trapnell, C., Pachter, L. & Salzberg, S.L. TopHat: discovering splice junctions with RNA-Seq. Bioinformatics 25, 1105–1111 (2009).

62. Trapnell, C., et al. Transcript assembly and quantification by RNA-Seq reveals unannotated transcripts and isoform switching during cell differentiation. Nat Biotechnol 28, 511–515 (2010).

